# Cuticle architecture and mechanical properties: a functional relationship delineated through correlated multimodal imaging

**DOI:** 10.1101/2022.12.19.521062

**Authors:** Nicolas Reynoud, Nathalie Geneix, Angelina D’Orlando, Johann Petit, Jeremie Mathurin, Ariane Deniset-Besseau, Didier Marion, Christophe Rothan, Marc Lahaye, Bénédicte Bakan

## Abstract

- Cuticle are multifunctional hydrophobic biocomposites that protect aerial organs of plants. Along plant development, plant cuticle must accommodate different mechanical constraints combining extensibility and stiffness, the corresponding structure-function relationships are unknown. Recent data showed a fine architectural tuning of the cuticle architecture and the corresponding chemical clusters along fruit development which raise the question of their impact on the mechanical properties of the cuticle.
- We investigated the in-depth nanomechanical properties of tomato fruit cuticle from early development to ripening, in relation to chemical and structural heterogeneities by developing a correlative multimodal imaging approach.
- Unprecedented sharps heterogeneities were evidenced with the highlighting of an in-depth mechanical gradient and a ‘soft’ central furrow that were maintained throughout the plant development despite the overall increase in elastic modulus. In addition, we demonstrated that these local mechanical areas are correlated to chemical and structural gradients.
- This study shed light on a fine tuning of mechanical properties of cuticle through the modulation of their architecture, providing new insight for our understanding of structure-function relationships of plant cuticle and for the design of biosinpired material.

## Introduction

Plant terrestrialization coincided with the development of the cuticle at the surface of aerial organs, to cope with harsh desiccating and UV-rich conditions (Niklas *et al*., 2017; Jiao *et al*., 2020) The cuticle fulfils multiple biological functions including the regulation of water and gas exchanges and the protection against environmental stresses (Martin & Rose, 2014; Fernández *et al*., 2021). Plant cuticle is also critical during the plant development as it prevents organ fusion (Sieber *et al*., 2000; Ingram & Nawrath, 2017; Renault *et al*., 2017) and provides a biomechanical support for the maintenance of the physical integrity of cuticle throughout the development and expansion of plant organs (Bargel & Neinhuis, 2005; Knoche & Lang, 2017).

Actually, the mechanical properties play a key role on the biological functions of cuticles, especially by supporting plant growth and resistance to environmental stress. Many studies have investigated these properties through tensile tests on isolated skins or cuticles (Petracek & Bukovac, 1995; Wiedemann & Neinhuis, 1998; Bargel & Neinhuis, 2004, 2005; Matas *et al*., 2004a,b; López-Casado *et al*., 2007; Domínguez *et al*., 2009; Lopez-Casado *et al*., 2010; Takahashi *et al*., 2012; Tsubaki *et al*., 2013; Khanal *et al*., 2013; Khanal & Knoche, 2014; Benítez *et al*., 2021). Because of its thick, astomatous and easy-to-isolate cuticle, the tomato (*Solanum lycopersicum*) fruit is a convenient model to study the mechanical properties of the cuticles at different scales. In facts, most of the available knowledge has been obtained on tomato fruit. The well-defined growth of tomato (Guillet *et al*., 2002; Renaudin *et al*., 2017) is also advantageous for studying cuticle stiffness and extensibility, the balance of which is necessary to avoid cuticle rupture in developing organs. In particular, the cuticle must accommodate the massive increase in volume during fruit expansion while ripening provokes the disassembly of cell walls, resulting in higher mechanical stress in the cuticle (Domínguez *et al*., 2009; Knoche & Lang, 2017; Jiang *et al*., 2019). Taken together, it appears that the mechanical properties of the cuticle depend primarily on the assembly and interactions of cuticular components, which evolve during organ development.

Indeed, the plant cuticle is a biocomposite made of a complex supramolecular assembly of monomeric and polymeric lipids, polysaccharides and phenolics (Fernández *et al*., 2016; Philippe *et al*., 2020; Reynoud *et al*., 2022). Cutin, the hydrophobic scaffold of cuticles is an insoluble polyester of oxygenated fatty acids (Hunneman & Eglinton, 1972; Bhanot *et al*., 2021) whose polymerization index can vary during the plant development (Philippe *et al*., 2016). The cutin matrix is further filled and coated by waxes (Busta & Jetter, 2018; Lee & Suh, 2022). Besides, polysaccharides are also entangled in the cuticle (CEP, cuticle-embedded polysaccharides). The CEP comprise highly methyl- and acetyl-esterified pectins, hemicelluloses and crystalline cellulose, which are structurally distinct from the non-cutinized cell wall polysaccharides (NCP). (Philippe *et al*., 2020). Finally, phenolic compounds including esterified hydroxycinnamic acids (Riley & Kolattukudy, 1975; Graça & Lamosa, 2010) and free and bound flavonoids are also found in the cutin polyester (Hunt & Baker, 1980; Reynoud *et al*., 2022). The composition and proportion of plant cuticle components vary during fruit growth and ripening (Domínguez *et al*., 2008; España *et al*., 2014b). Recently, fine chemical Raman mapping of the tomato fruit cuticle revealed the in-depth spatial heterogeneity of its components (lipids, polysaccharides, phenolics) (González Moreno *et al*., 2022) including the cutin polymer matrix (Reynoud *et al*., 2022). The additional observation of the fine architectural tuning of the corresponding chemical clusters along fruit development raise the question of their impact on the mechanical properties of the cuticle. To gain further insights into these relationships, detailed information, at the nanoscale level, on the mechanical properties of the cuticle and their changes in the developing fruit are required.

To assess the in-depth mechanical properties of the Cutin Polymer Matrix (CPM) over cherry tomato fruit development in relation to chemical and structural heterogeneities, we designed a correlated multimodal imaging methodology. Atomic Force Microscopy (AFM) provides the sensitivity to address the mechanical properties of tissues at the nanometric scale (Dufrêne *et al*., 2017). This technique has been successful in depicting the mechanical heterogeneity of a wide range of biological tissues including cell wall and synthetic polymers (Xi *et al*., 2015; Melelli *et al*., 2020; Stoica *et al*., 2021). AFM was also used to probe the surface of tomato cuticle (Round *et al*., 2000; Isaacson *et al*., 2009). Focusing on the polymeric structure of the plant cuticle, we combined Peak-Force Quantitative Nanomapping (PF-QNM) mode of AFM with hyperspectral imaging techniques such as confocal Raman microspectroscopy and Optical PhotoThermal InfraRed (OPTIR) (Zhang et al., 2016). These approaches revealed that the CPM exhibits distinct in-depth mechanical areas with specific dynamics during the phases of cell expansion and fruit ripening that are correlated with chemical and structural gradients.

## Material and methods

### Sample preparation

Cherry tomato (*S. lycopersicum* var. cerasiforme WVa106) plants were grown in controlled glasshouses as previously described (Alhagdow *et al*., 2007). Flowers were tagged at anthesis and fruits at 10, 15, 20, 25, 30, 35 and 40 Days Post Anthesis (DPA) were harvested. Exocarp was collected at the equatorial part of the fruits, chemically fixed and impregnated in paraffin as previously described (Reynoud et al., 2022). Paraffin blocks were cut with an ultramicrotome (Leica EM UC7, Leica Microsystems SAS, Nanterre, France) equipped with diamond knifes (Histo and Ultra, Diatome, Nidau, Switzerland) to obtain 1µm-thick cross-sections. Sections were placed on BaF_2_ windows and paraffin was removed with successive baths of methylcyclohexane, ethanol and water. In addition, to study the susceptibility to cutinase of the cutin polymer matrix (CPM), 1µm-thick cross sections for 20 and 35 DPA stages were subjected to cutinase (Humicola insulens NZ 51032, Chiral Vision) treatment for 4h at 37°C as previously described (Philippe *et al*., 2020).

In parallel, tomato fruits were peeled off and skins prepared as isolated cuticle as previously described (Philippe *et al*., 2016). Cutin samples were obtained after removing waxes and non-bound phenolic with chloroforme:methanol (2/1, v/v).

### Correlated Multimodal Imaging

Correlated multimodal imaging was performed on the same sample per developmental stages and measurements were conducted in controlled environment (ambient air pressure and temperature of 20-25°C).

#### Atomic Force Microscopy (AFM)

Measurements of the tomato CPM nanomechanical properties were performed on a multimode 8 atomic force microscope (Bruker Nano Surface, Santa Barbara, CA, USA) equipped with RTESPA-150 (Bruker AFM Probes, Camarillo, CA, USA) and operated in Peak-Force Quantitative Nanoscale Mechanical mapping (PF-QNM). A deflection sensitivity of 12 nm.v^-1^ was determined on a sapphire surface (12M PF-QNM, Bruker AFM Probes, Camarillo, CA, USA) with an elastic modulus of 400GPa. The spring constant was determined after the thermal tune procedure using the Sader’s methods (https://sadermethod.org/; Sader et al., 1995). Before and after acquisition of a set of samples, calibrations of tips radius were evaluated using PS-LMDE-12M (Bruker AFM Probes, Camarillo, CA, USA), i.e., a blend of PolyStyrene (PS) with an elastic modulus of 2 GPa and Low-Density Polyolefin Elastomer (LMDE) with a module of elasticy of 0.16 GPa. The spring constant ranged between 10 and 13 N/m and the tip radius between 8 and 10 nm for the probes used in this study. Nanomechanical imaging was performed with a Peak force setpoint set at 6nN, and an oscillation of 160kHz and 40µm/s for scan rate. Hertz’s Model (Hertz, 1882) was chosen over the Derjaguin-Muller-Toporov (DMT) (Derjaguin *et al*., 1975) to process force-distance curve as the fit was better and adhesion forces were found to be quite homogeneous in our samples. Moreover, the indentation applied was below 1% of the sample height, meeting the rule for the Hertz model (Zdunek & Kurenda, 2013). Several acquisitions per sample were done on 1µm-thick cross-sections and at the surface of isolated cutin sample with at least two images per developmental stage processed with Gwyddion software (http://gwyddion.net; Nečas and Klapetek, 2012). Topographic images were levelled by least square method and mechanical measurements such as profiles or sampling in CPM areas, were performed on filtered elastic modulus maps to remove outliers, i.e., elastic modulus higher than the upper limit of the AFM probes were discarded. After assessing normality and homoscedasticity, mean comparison were achieved through two-way parametric ANOVA and further evaluated through Tukey HSD using R software (www.r-project.org) and “FactoMineR” package (Lê *et al*., 2008).

#### Confocal Raman microspectroscopy

Raman imaging was performed with an inVia™ Renishaw confocal Raman microspectrophotometer and operated as previously described (Reynoud et al., 2022) with minor adjustments. Maps were recorded on the same area as acquired with AFM, with spatial resolutions of 0.5 µm in both x- and y-directions. Cosmic rays were removed from Raman spectra using the WiRE 4.23 software (Renishaw, UK). To improve the signal to noise ratio and the specificity of Raman signals, an Extended Multiplicative Scattered Correction (EMSC) method (Kerr & Hennelly, 2016) combined with Principal Components Analysis (PCA) noise filtering PCA (He *et al*., 2020) was used. Preprocessing was executed in Quasar software (version 1.5.0, https://quasar.codes) and spectra further analysed in R software.

In parallel, molecular orientations within the CPM were investigated at 20 and 40 DPA stages as previously described (Reynoud *et al*., 2022) by acquiring the same area with distinct laser polarization directions: one set parallel to the plane of incidence (0°) and the other one set orthogonal to the plane of incidence (90°).

#### Optical PhotoThermal InfraRed (OPTIR)

OPTIR imaging was performed on a mIRage™ Infrared microscope (Photothermal Spectroscopy Corp., Santa Barbara, CA, USA). Samples, i.e., 1µm-thick cross-sections on BaF_2_ window (see ‘Sample preparation’ subsection), at 20, 30 and 40 DPA stage were placed on the motorized plate and the same area recorded in Raman and AFM was mapped. The IR source was a pulsed, tunable four-stage QCL device, scanning from 950 to 1950 cm^-1^, with the power set to 13% and duty cycle of 1%. The probe laser was a visible laser at 532 nm with the power set at 5.7%. Before measurement, calibration was performed on carbon black reference to optimize laser position for 1850, 1550, 1300 and 1050 cm^-1^, each wavenumber corresponds to one chip of the QCL device. Mapping was performed with a 40× objective (Schwarzschild, NA = 0.78) achieving a spatial resolution of around 0.5 µm in both x and y directions with spectral data points spaced by 2 cm^-1^. Each data point is an average of four spectra.

### Correlative analysis

In order to associate mechanical properties with chemical information, Raman or OPTIR maps were first subjected to pixel co-registration to spatially colocalize chemical data with mechanical data from elastic modulus maps. The white light images of the area recorded in Raman or OPTIR were superimposed onto AFM topographic maps using at least 10 visual features as anchoring points. These linear transformations were performed in Mountains Map 9 (Digital Surf, Besançon, France) software and newly spatialized mechanical maps were extracted to be further processed in R software. As AFM and Raman or OPTIR imaging have distinct spatial resolution, degradation of AFM resolution by averaging was perform with “raster” (https://cran.r-project.org/web/packages/raster/) and “terra” (https://cran.r-project.org/web/packages/terra/) packages to fit the 0.5 µm²-size of pixel in Raman and OPTIR. Inter-modality Pearson correlations between hyperspectral data and mechanical properties were conducted using R package “Hmisc” (https://cran.r-project.org/web/packages/Hmisc/index.html).

## Results

### The mechanical properties of the tomato cutin polymer matrix display spatiotemporal heterogeneities

To assess the in-depth mechanical properties of the cutin polymer matrix of the tomato fruit from early developmental stages (10 DPA) to ripening (40 DPA), crosssections of isolated exocarps were imaged by AFM with PF-QNM mapping mode. First, to validate our experimental set-up, we compared at the same development stages and with identical AFM conditions, the mechanical properties, i) at the surface of isolated non-fixed tomato fruit cutin (Fig. **1a**) and ii) on the outer edge of the fixed cutin cross-sections (Fig. **1b**). Young’s modulus of the surface was assessed on cuticular ridges between two depressions (Fig. **1a**, square), which represent epidermal cell boundaries (Isaacson *et al*., 2009). From 15 DPA to 40 DPA a significant increase of the Young’s modulus, from 145 ± 52 to 542 ± 302 MPa, was observed. These results perfectly fit previous data obtained from isolated tomato cuticles during the fruit growth and ripening (Andrews *et al*., 2002; Bargel & Neinhuis, 2005; Benítez *et al*., 2021). Likewise, from 15 to 40 DPA, Young’s modulus at the edge of cuticle cross-sections significantly increased from 139 ± 115 to 697 ± 415 MPa (Fig. **1c**). Despite these differences between absolute values, a similar increase was observed with a positive correlation (0.45, p-value < 0.001). Therefore, we further imaged the tomato fruit cross-sections from 10 to 40 DPA (Fig. **2**).

**Fig. 1.**
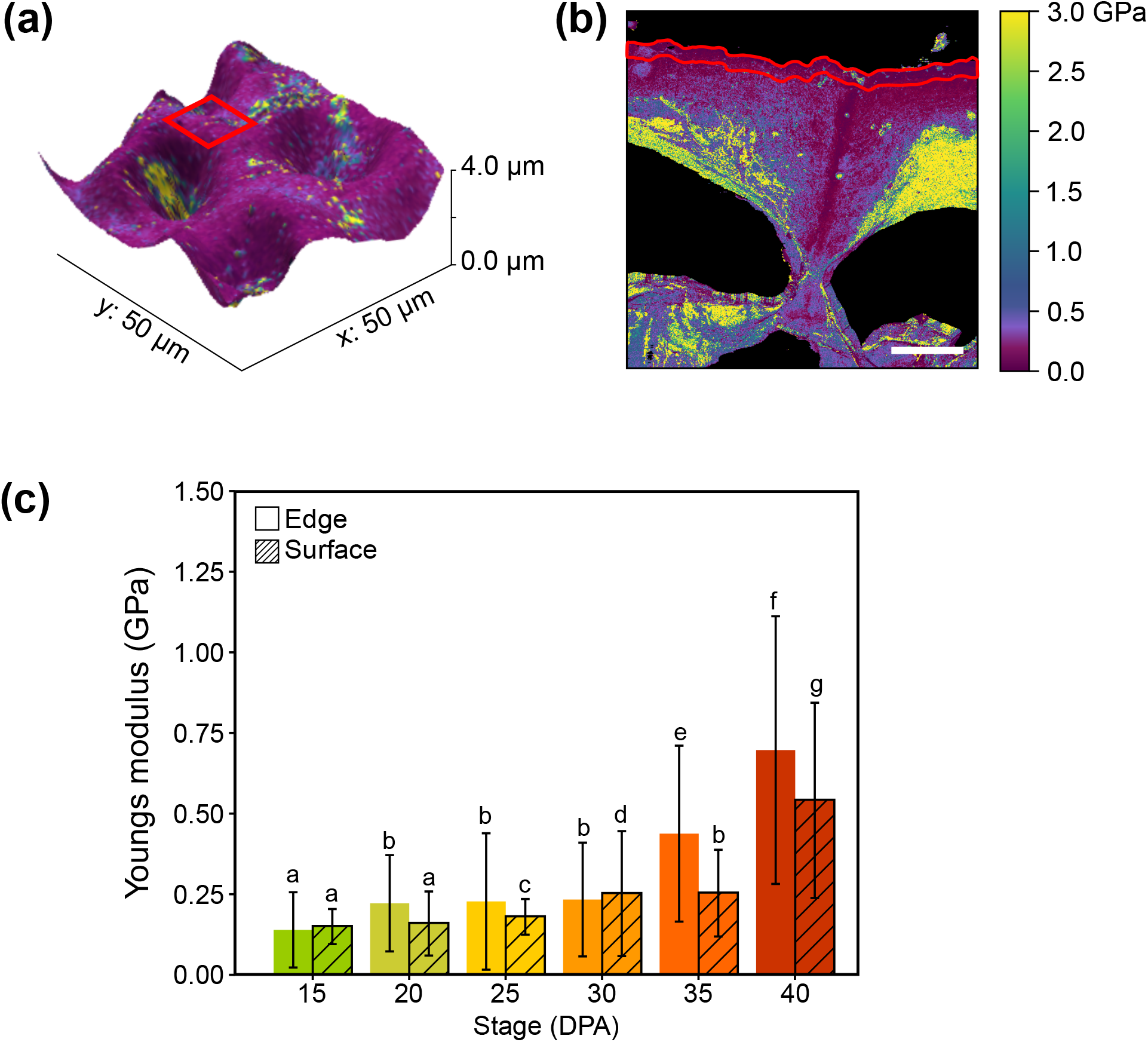
Mechanical properties of the surface of isolated cuticles and at the edge of cuticle cross-sections along tomato fruit development. (a) Illustration of elastic modulus map superimposed onto the volumetric topography of isolated and non-fixed tomato cuticle at 20 Days Post Anthesis (DPA) stage. Red square represents a window of 7x7 µm^2^. (b) Illustration of elastic modulus map of a fixed cutin crosssection at 20 DPA. The red polygon delimits an area of 0.5 µm-width on the entire length of the cross section. Bar, 5 µm. (c) Comparison of mechanical properties on the surface and at the edge of the cutin polymer matrix at different developmental stages of tomato fruit. The letters above error bars indicate a significant difference between developmental stages and sampled regions (P < 0.05, ANOVA and Tukey HSD).

**Fig. 2.**
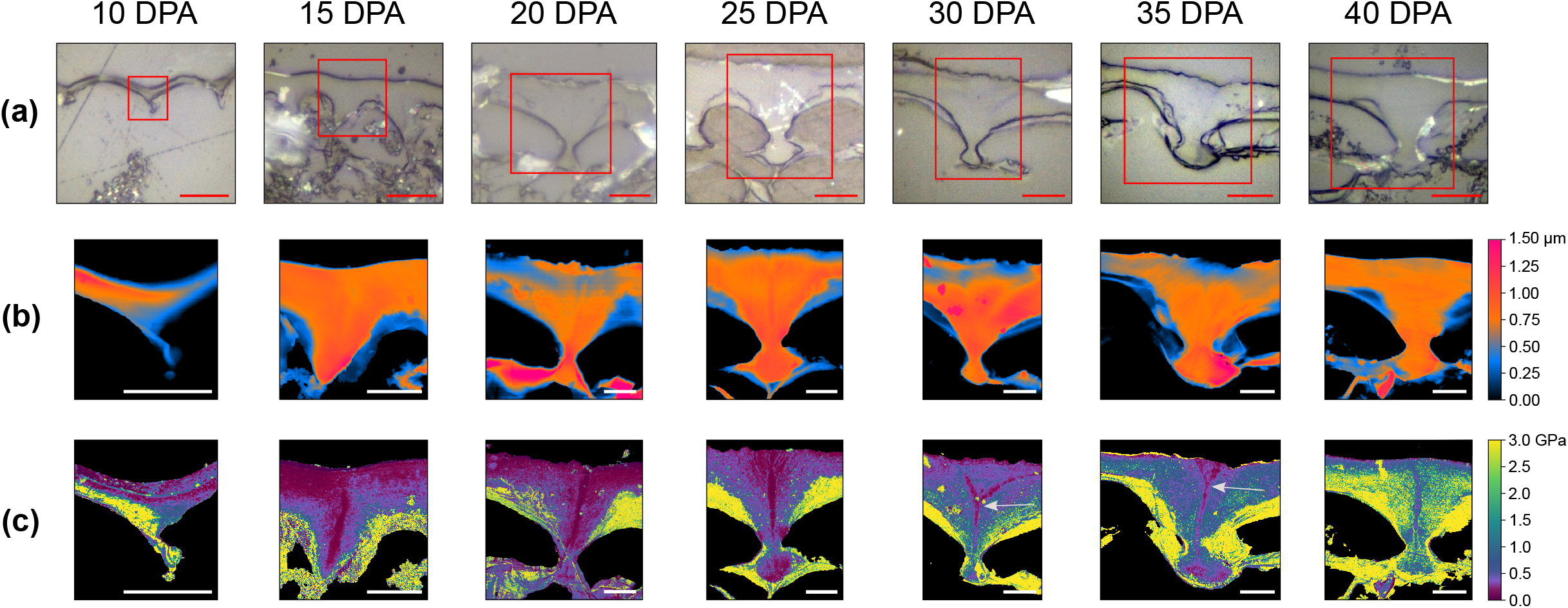
Peak-Force Quantitative Nanoscale Mechanical (PF-QNM) mapping of the cutin polymer matrix during tomato fruit development. Two maps per developmental stage were acquired, representative images are shown. (a) Light visible images of cuticles with red rectangles highlighting areas that were investigated with PF-QNM. (b) Topography. (c) Young’s modulus. DPA, Days Post Anthesis. Bars represent 10 µm in (a) and 5 µm in (b) and (c). White arrows indicate the division of the central furrow at 30 and 35 DPA.

Unprecedented sharp differences in the Young’s modulus within the CPM were highlighted (Fig. **2c**). For instance, a gradient in the elastic modulus was observed from the external part of the cuticle to the cell wall. In addition, a clear central furrow with a lower Young’s modulus was evidenced during fruit growth, from 15 DPA to 40 DPA (Fig. **2c**). The absence of a central furrow at 10 DPA suggests that it is formed later, concomitantly with the cuticular peg (Segado *et al*., 2016) and onset of rapid deposition of cutin in the cuticle (Petit *et al*., 2014). At the 30-35 DPA stage, the furrow did not span the entire in-depth of the CPM, but divided near the outermost part of the cuticle facing the cuticular pegs (see arrows Fig **2c**). Besides, during tomato fruit development, Young’s modulus of the CPM was modified although not homogeneously (Fig. **2c**). Accordingly, we further assessed the specific spatiotemporal heterogeneities of elastic modulus in the cutin polymer matrix focusing on these specific areas.

### Spatiotemporal dynamics of the mechanical properties of CPM

The CPM mechanical properties were transversally and longitudinally measured during fruit development to assess their spatiotemporal dynamic (Fig. **3**).

**Fig. 3.**
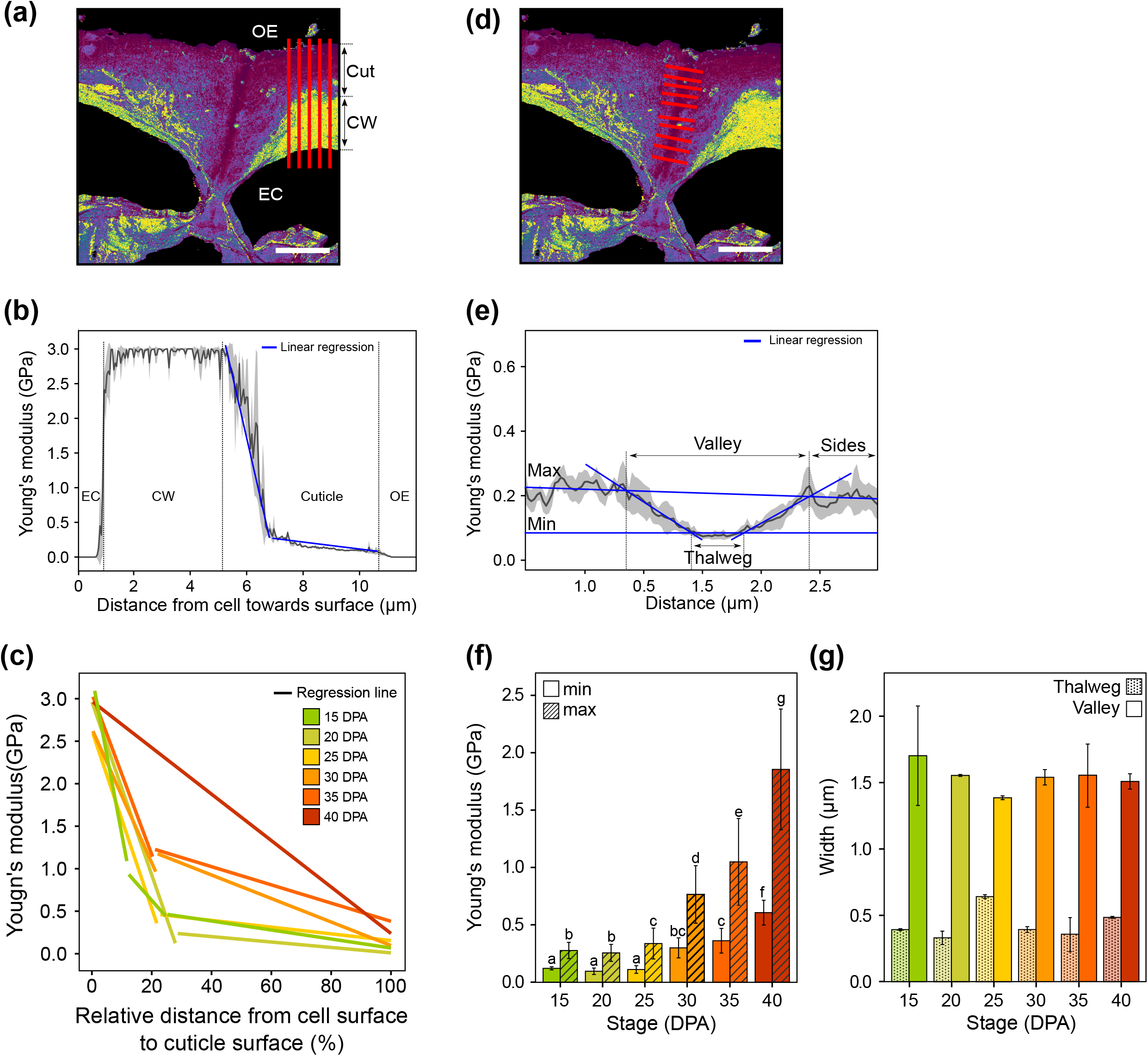
Spatiotemporal dynamics in mechanical properties of region of the cutin polymer matrix (CPM). (a) Illustration of in-depth mechanical profile sampling (red lines) of the CPM at 20 days post anthesis (DPA) from the surface of epidermal cell (EC) to the surface of the cuticle (OE). Cut, Cuticle; CW, cell wall. Each profile consisted of 160 to 235 equally interspersed points, according to the in-depth cuticle thickness, with each point being a mean of 8 measurements. Two maps per developmental stage were acquired with 7 sampling lines per stage. (b) Illustration of linear regressions (blue lines) performed on the decreasing phase of the average elastic modulus extracted from the in-depths profiles of a representative map at 20 DPA. Data are expressed as mean (solid lines) ± SD (shadowed area). To visualize the complete profiles, readers are referred to Fig. S1. (c) Linear regression curves of the in-depth elastic modulus of the CPM, according to the tomato developmental stage. For proper comparison, the distance from the cell surface to the cuticle surface was normalized and expressed as percentage. (d) Illustration of mechanical profile sampling (red lines) of the central furrow of the CPM at 20 DPA. Ten profiles per developmental stage were sampled with each profile consisting of 120 equally interspersed points over 3µm; each point being a mean of 8 measurements. (e) Illustration of linear regressions (blue lines) performed on the average elastic modulus extracted from the transverse profile of a representative map at 20 DPA. Data are expressed as mean (solid lines) ± SD (shadowed area). Regressions were calculated on the minimum and maximum plateau and on both sides of the depression. Size of the furrow was estimated on the upper part of the curve, between 1^st^ and the 4^th^ bending points of the modulus curve (‘Valley’) and on the lower part of the curve, between the 2^nd^ and the 3^rd^ bending points (‘Thalweg’). To visualize the complete profiles, readers are referred to Fig. S2. (f) Dynamics of elastic modulus of the central furrow and besides, over the developmental stage of tomato fruit. Values are mean of two maps acquired per developmental stages and calculated on the ‘Thalweg’ (min) and on the ‘Sides’ (max). Letters above error bars indicate significant difference (ANOVA, P <0.05 and Tukey HSD). (g) Dynamics of the central furrow size (‘Thalweg’, ‘Valley’) of the CPM over tomato fruit development. Values are mean ± SD (n=2). Bars, 5µm.

The in-depth mapping of the elastic modulus was done from the surface of the epidermal cells to the surface of the cuticle, on the periclinal part of the CPM to avoid any biases due to thickness heterogeneities of cell wall and cuticle (Fig. **3a**). Linear regressions were performed (Fig. **3b** and **S1**) and a progressive increase of the CPM elastic modulus was evidenced during development, but with distinct profiles (Fig. **3c**). At 15 DPA, the mechanical gradient showed three phases, with two apparent inflection points. From 15 to 25 DPA, and from 30 to 35 DPA, two phases were observed with distinct slopes whereas, at 40 DPA, a single linear increase of the elastic modulus was observed.

Likewise, we looked through to Young’s modulus profiles of the furrow within the CPM at the different developmental stages (Fig. **3d**). Linear regressions were performed on the maximum, the minimum and on both sides of the slope (Fig. **3e** and **S2**) to compare mechanical properties (Fig. **3f**) and size of the furrow over fruit development (Fig. **3g**). The size of the furrow was evaluated according to Young’s modulus value. Two different zones were highlighted within this furrow, i.e., the central soft area (expressed as ‘Thalweg’) and the overall furrow size (hereafter named ‘Valley’) (Fig. **3e**). A significant increase of Young’s modulus was measured in both the Thalweg (Fig. **3f**, min) and the ‘hard’ sides (Fig. **3f**, max) of the furrow. Indeed, between 15 and 40 DPA, the elastic modulus increased from 122 ± 8 to 605 ± 72 MPa for the central furrow and from 276 ± 3 to 1842 ± 77 MPa for the sides of the furrow. Interestingly, during fruit expansion, i.e., from 15 to 25 DPA, the difference between the elastic modulus of the ‘Thalweg’ and the Sides’ of the furrow were almost constant and started to increase between 25 and 30 DPA, i.e., at the end of rapid fruit expansion and onset of ripening (Guillet *et al*., 2002). The size of this central furrow was monitored during tomato fruit development. The overall size of the furrow (‘Valley’) was quite homogeneous (around 1.5 µm) during development. Conversely, the central part of the furrow (‘Thalweg’) was around 0.38 µm for all developmental stages except for the 25 DPA which showed a wider Thalweg of 0.63 µm (Fig. **3g**). At this stage, the difference between Valley and Thalweg sizes was lower than that at other developmental stages indicating a more abrupt transition from high-to-low elastic modulus in the furrow.

Taken together, these peculiar spatiotemporal dynamics of the nanomechanical properties of the cutin matrix during fruit development led us to look for possible relationships with the chemical composition of these areas.

### Chemical gradients and in-depth mechanical heterogeneities are related

To investigate these in-depth mechanical heterogeneities both Raman and Optical PhotoThermal InfraRed (OPTIR) spectroscopies were used. Cluster analyses from the OPTIR maps mainly revealed a gradient in the cutin/polysaccharides ratio from the cuticle surface toward the surface of epidermal cells (Fig. **4a** and **S3**) (Philippe *et al*., 2020). Indeed, as already indicated from macroscopic analyses (isolated peels or cuticle), the amount of embedded polysaccharides has a significant impact on the elastic modulus (López-Casado *et al*., 2007). In this regard, Raman mapping provides a finer chemical clustering of the CPM, i.e., lipids, polysaccharides and phenolic compounds (Fig. **4a** and **S3**) (Reynoud *et al*., 2022). From these Raman maps, the radial profile of specific bands of CPM components were obtained (Fig. **4b** and **S4**). Then, Pearson correlations were calculated between elastic modulus and relative Raman intensity of these specific bands (Fig. **4b**, bold values in panels). Statistically significant correlations were obtained at every developmental stage, although the correlation values were lower at 35 and 40 DPA as a result of the complexification of CPM with increased numbers of chemical clusters (Reynoud *et al*., 2022). Positive correlations were observed between the elastic modulus and the Raman intensity of crystalline cellulose (e.g., 0.76 at 30 DPA) and pectins (e.g., 0.75 at 30 DPA) (Fig. **4b**). The same conclusion could be drawn using a generic band for polysaccharides (1375 cm^-1^), due to the HCC, HCO and COH bending deformations (Chylińska *et al*., 2014). At the opposite, negative correlations were evidenced with the Raman intensity of bands assigned to lipids (e.g., -0.70 at 20 DPA) and p-coumaric acid (pCA) (e.g., -0.69 at 20 DPA) (Fig. **4b**). Upon ripening, CPM accumulates phenolics as evidenced by a generic band for phenolics (1170 cm^-1^) that is assigned to δ(CH) of aromatic rings (Świsłocka *et al*., 2012), related to phenolic acids and flavonoids.

**Fig. 4.**
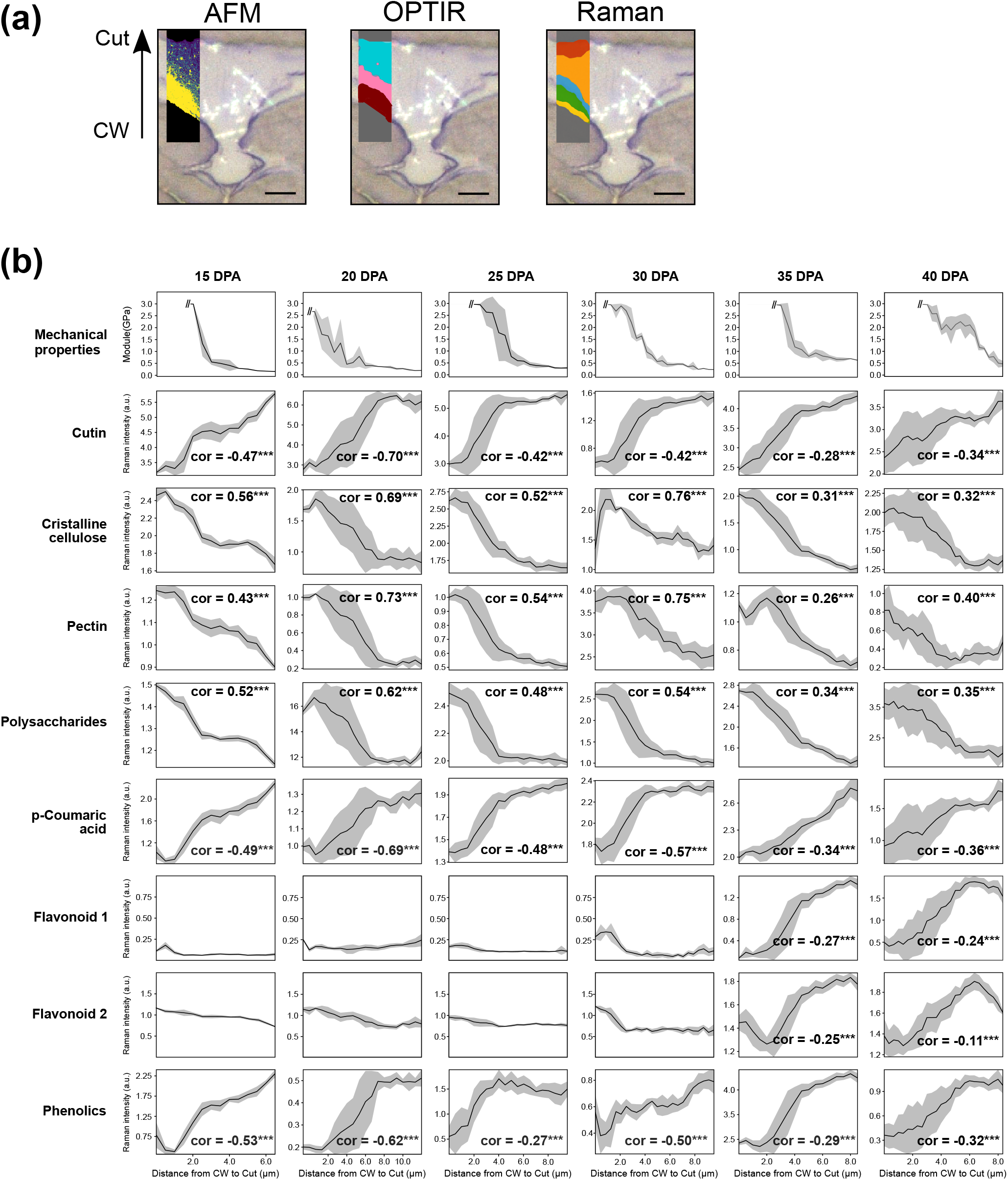
Chemical profile of the cutin polymer matrix (CPM) over the in-depth thickness. (a) Illustrations of sampling zone in cross-section of tomato cuticle at 25 DPA for AFM maps (left), OPTIR (middle) and Raman (right). For both OPTIR and Raman data, the sampled zone is represented by a k-means clustering map with three clusters for OPTIR data and five clusters for Raman. To see the mean spectra associated to the different clusters, readers are referred to Fig. S3. Bars, 5µm. (b) Chemical composition assessed by Raman spectroscopy and elastic mechanical properties determined by PF-QNM from the cell surface to the cuticle surface of the CPM over tomato fruit development. Chemical profiles of CPM components were obtained on area-mean normalized data to avoid biases due to difference in crosssection thickness (see Fig. 2) and expressed as Raman intensity with arbitrary units (a.u.). For each component a specific band was selected according to a homemade reference database (Reynoud et al., 2022): Cutin (1440 cm^-1^), Crystalline cellulose (380 cm^-1^), Pectin (852 cm^-1^), Polysaccharides (1375 cm^-1^), p-Coumaric acid (1605 cm^-1^), Flavonoid 1 (547 cm^-1^), Flavonoid 2 (1550 cm^-1^) and Phenolics (1170 cm^-1^). Bold values in charts represent Pearson correlations (*P value < 0.05; ** P value < 0.01; *** P value < 0.001) between elastic modulus and Raman intensity. See Fig. S4 for full spectra.

Flavonoids are specifically accumulated in the cuticle during ripening (Laguna *et al*., 1999; España *et al*., 2014b) including a fraction associated to CPM (Fig. **4b**) (Hunt & Baker, 1980; Domínguez *et al*., 2009). Interestingly, negative correlations of the intensity of both flavonoid specific bands with elastic modulus were highlighted. Accordingly, at the submicron scale, our results indicate that the cutin-associated flavonoids do not account for the modification of Young’s modulus related to flavonoid accumulation of the isolated cuticles (España *et al*., 2014a). Together, these results indicate that the nanomechanical heterogeneities revealed in the CPM are correlated to chemical gradients.

### The central furrow displays a fine structural arrangement and progressive chemical inhomogeneity

To decipher the specific mechanical properties of the central furrow and its relationships with the chemical composition and structure of the CPM, a multimodal approach was conducted combining Raman imaging and Infrared mapping at three developmental stages, i.e., 20, 30 and 40 DPA. Two areas of the CPM were compared, the central furrow and the sides of the furrow (Fig. **3e** and **S5**). At 20 and 30 DPA, principal component analysis (PCA) from Raman data showed no strong specific spectral fingerprints of the central furrow until the 40 DPA stage whereas it was more progressive for OPTIR data (Fig. **5a**). PCA loadings of OPTIR data (Fig. **S6a**) evidenced that the central furrow was rather composed of cutin whereas polysaccharides were progressively excluded to the sides. Loading of Raman data (Fig. **S6b**) indicated a change of the relative accumulation of phenolic compounds in the furrow sides along with crystalline cellulose whereas pectins concentrated in the central furrow. These multimodal spectroscopic approaches indicated that the central furrow adopts a slight but distinct chemical composition than the furrow sides along tomato fruit development.

**Fig. 5.**
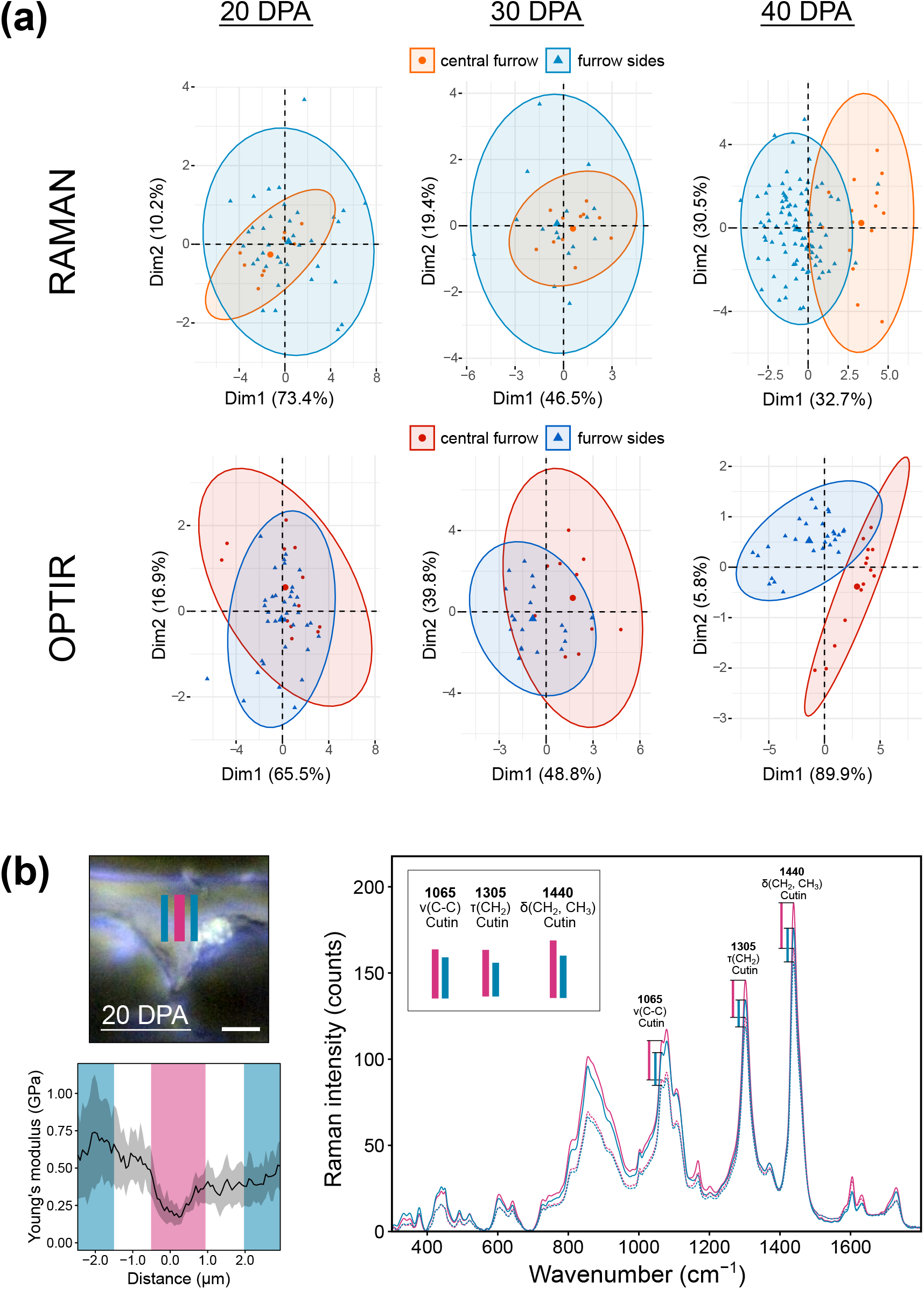
Chemical composition and macromolecular arrangement of the central furrow over the development of the tomato CPM. (a) Principal component analysis (PCA) of Raman (up) and OPTIR (down) datasets of the central furrow compared to furrow sides at 20, 30 and 40 days post anthesis (DPA). Sampling zones and PCA loadings are found in supplemental information (Fig. S5 and S6). (b) Macromolecular orientation of lipids within the central furrow compared to the furrow sides at 20 DPA. The laser polarization direction was modified from parallel (0°) to orthogonal (90°) to the cuticle surface. For each polarization, two maps were acquired at 20 and 40 DPA (Fig S7). The light image (upper left) represents the area of the cuticle that were sampled to assess the molecular orientations within the central furrow (pink) and the furrow sides (blue) with the corresponding elastic modulus (lower left). Bar, 5µm. Data are expressed as mean (solid lines) ± SD (shadowed area). For both central furrow and furrow sides, mean Raman spectra for each polarization direction were calculated (right). The parallel (solid lines) to orthogonal (dashed lines) ratio of intensity was illustrated by vertical bars for three bands related to lipids: two bands attributed to CH_2_ deformations, i.e., τ(CH_2_) at 1305 cm^-1^ and δ(CH_2_,CH_3_) at 1440 cm^-1^ and one band assigned to ν(C-C) vibration at 1065 cm^-1^. Insets represents the lined up different intensity ratio (bars) for proper comparison between cuticle areas.

Then, the specific molecular orientation of CPM components was further assessed by Raman spectroscopy using a linear polarized laser. By following changes in band intensity due to different settings of polarization, parallel (0°) and orthogonal (90°) to the cuticle surface, specific molecular orientations could be inferred (Fig. **5b**). We focused on lipid bands, as the central furrow was mostly composed of cutin. Two bands related to CH_2_ modes of vibrations: δ(CH_2_, CH_3_) and τ(CH_2_), and one related to C-C stretching vibrations (Czamara *et al*., 2015) were found to changes according to polarization (Fig. **5b**). A higher intensity with the 0° polarization direction suggested a preferential orientation of lipids parallel to the cuticle surface for both 20 and 40 DPA stages (Fig. **5b** and **S7**). Differences between the central furrow and its sides were observed at both stages (Fig. **5b** and **S7**) illustrating a distinct macromolecular arrangement of lipids of these areas.

To further gain insight into the specific chemical composition and structure of the central furrow, the susceptibility to cutinase was checked. AFM experiments were conducted on this furrow area enabling AFM topography. Upon cutinase treatment, the central furrow was sharply carved out compared to the furrow sides at 20 DPA and 35 DPA as well (Fig. **6**) This acute difference of cutinase susceptibility within the central furrow could be related to higher accessibility of the enzyme, in agreement with the difference in macromolecular arrangement highlighted by Raman mapping (Fig. **5b**). Moreover, cutinase is more active towards primary than secondary ester bounds (Lin & Kolattukudy, 1978). Accordingly, the significant impact of cutinase on the furrow topography could be due to a lower reticulation degree of the cutin polyester (i.e., with less secondary ester bonds) in the furrow than on the sides of the furrow.

**Fig. 6.**
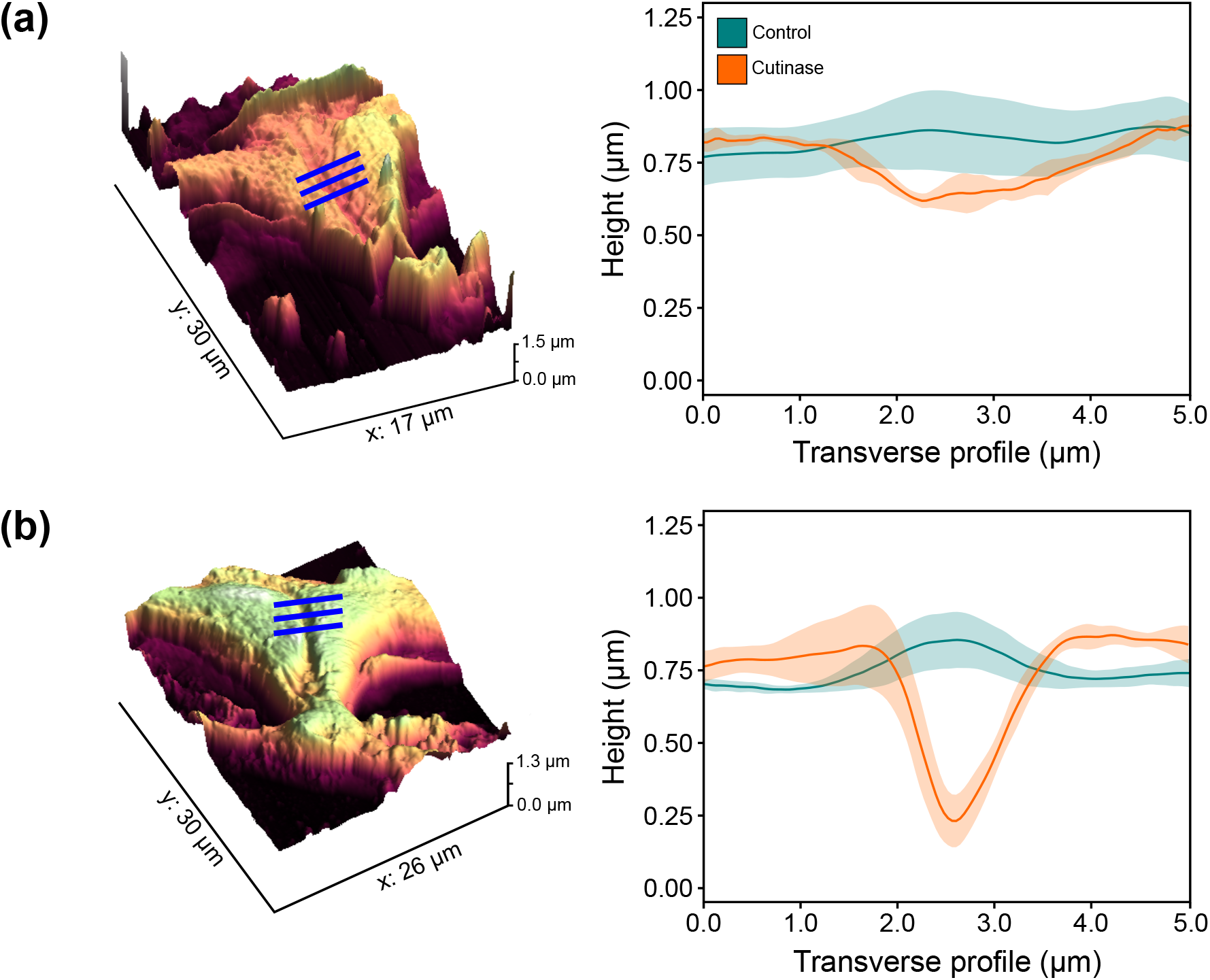
Susceptibility of the cutin polymer matrix (CPM) to cutinase hydrolysis. Topography of the CPM either subjected to cutinase treatment or not at 20 (a) and 35 days post anthesis (DPA). The 3D topographies (left) illustrate the carving effect of the cutinase in the central furrow with the blue line representing the sampled transverse profiles within the CPM. The corresponding topographic profiles (right) were calculated as a mean of 10 profiles over 3 µm, each profile being a mean of 8 measurements. Data are expressed as mean (solid lines) ± SD (shadowed area). Two maps per developmental stages were acquired.

Together our results showed variations in nanomechanical properties in the central furrow that are associated not only with chemical modifications but also with macromolecular arrangements within the CPM.

## Discussion

### Heterogeneity of local mechanical properties of the cutin polymer matrix: a way to address challenges during fruit growth?

Elucidation of relationships between the architecture of the cutin-polysaccharide assemblies and the functional properties of the cuticle is critical to understand and modulate plant cuticle functionalities. All along fruit development, the mechanical properties of the cuticle must adapt to the trade-off between fruit expansion and maintenance of cuticle integrity (Knoche & Lang, 2017).

In the present study, in depth nanomechanical mapping of the cutin polymer matrix evidenced unprecedented heterogeneities in the cuticle, including “soft” (exhibiting elastic modulus around 300-500 MPa) and “hard” (modulus around 1.5-2.5 GPa) domains. Such spatial nanomechanical heterogeneity, maintained throughout the development, was not observed at the surface of isolated cuticle (Fig. **1**) (Round *et al*., 2000; Isaacson *et al*., 2009). Most strikingly, a “central furrow” displaying a lower elastic modulus was evidenced in the CPM between the anticlinal walls of adjacent epidermal cells (Fig. **2**). The biological function of this peculiar ‘soft’ central furrow maintained during the fruit expansion phase has to be addressed. The lower Young’s modulus suggests a higher mechanical compliance (Rusin & Kojs, 2011). A similar situation is observed with some Algae where specialized soft and extensible tissue (geniculum) are inserted between stiff and rigid calcified cell wall (intergeniculum) (Denny and King 2016). In addition, this central region of plant cuticle is currently described as a place for the deposition of cuticle material to form the so-called ‘cuticular pegs’ (Deas & Holloway, 1977). From an anatomical point of view, the ‘central furrow’ could be compared to the ‘middle lamella’ in-between two primary cell walls. In accordance with this hypothesis, this area is enriched in pectins and poor in cellulose (Fig. **5a** and **S6**) as observed in the ‘middle lamella’ (Jarvis *et al*., 2003). In this regard, middle lamella in plant cell wall acts not only in cellular adhesion but also plays a role in the mechanical resistance under tension or compression (Zamil & Geitmann, 2017). Previous studies suggest that the cuticle above the anticlinal peg concentrate the mechanical stress (Knoche & Lang, 2017). Accordingly, the presence of a ‘soft’ area might play a key role in the biomechanical properties of the epidermis by “absorbing” mechanical stress, while allowing tissue to extend. In line with this hypothesis, the size of the central furrow (‘Thalweg’) is maximum at 25 DPA corresponding to the maxima of fruit expansion rate (Guillet *et al*., 2002). Interestingly, at a macroscopic scale, tensile test of isolated cuticle highlighted a higher extensibility at this development stage of the tomato fruit (España *et al*., 2014b; Benítez *et al*., 2021). These mechanical features are probably associated to biochemical modification within the CPM. Indeed, the 25 DPA stage was also highlighted as a turning point in macromolecular rearrangement of polysaccharides embedded in the cuticle (Reynoud *et al*., 2022). Furthermore, our data indicate that the lipid polyester of this central ‘soft’ area has also peculiar structural features including a specific lipid orientation revealed by Raman mapping and higher sensitivity to cutinase hydrolyses (Fig. **5** and **6**), suggesting a specific macromolecular organization and a lower reticulation degree of the polyester. Interestingly, in cutin-inspired polyesters, the lower polymerization index was also associated with a lower elastic modulus and higher extensibility (Marc *et al*., 2021). Altogether, our study highlighted a specific design of tomato cuticles architectures leading to alternating soft and hard interfaces. Plant cuticle is therefore another example of biological material featured by site specific mechanical properties to address environmental or developmental constraints (Liu *et al*., 2017).

### The cutin-polysaccharide continuum: toward the design of a structural gradient during fruit development

The cutin-polysaccharide continuum plays a pivotal role in the mechanical properties of the plant cuticle. Indeed, from tensile tests of either fruit skin or isolated cuticles, it was suggested that the extent of cell-wall cutinization, is responsible for the higher mechanical stiffness in a crack resistant tomato cultivar (Matas et al., 2004a). Likewise, the proportion of polysaccharides within the cuticle was positively correlated to the elastic modulus (López-Casado *et al*., 2007; Takahashi *et al*., 2012), although the contribution of each component to cuticle mechanical properties is difficult to determine.

In the present study, correlated multimodal imaging enabled new insight in the cutin-polysaccharide continuum and demonstrated a fine tuning of its nanomechanical properties during fruit development. From fruit expansion to fruit maturation phases, an overall increase of the elastic modulus (up to 6 fold) was observed. Furthermore combining the hyperspectral and nanomechanical mapping, the impact of the cutin-polysaccharide ratio on the mechanical properties of CPM agrees with the strain-hardening behavior provided by tensile tests measurements of isolated cuticles (Matas *et al*., 2004a; Bargel & Neinhuis, 2005; Benítez *et al*., 2021). More precisely, our study highlighted three types of interface, i.e., transition zones, in the cutin-polysaccharide continuum (Fig. **3c** and **S1**) that coincides with sharply contrasted phases of tomato fruit development, i.e., the fruit expansion (10-25 DPA) and the ripening process (30-35 DPA) leading to the red ripe stage (40 DPA) (Guillet *et al*., 2002; Renaudin *et al*., 2017). From 10 DPA to 30 DPA, in-depth elastic modulus mapping highlighted a sharp interfacial zone between the cutin-rich and the polysaccharides-rich areas. During maturation, a gradual broadening of this interfacial zone turned into a continuous gradient transition (Fig. **3c**). In composites, structural transitional mechanical gradients are currently considered as a way to accommodate mechanical properties mismatches (e.g. elastic modulus) by a smooth transition and to provide stress relief at the interface between dissimilar materials (i.e., lipid polyester and cell wall polysaccharides) (Liu *et al*., 2017). Such structural gradients result in an increased toughness and have been observed in many biological materials such as the dentin-enamel junction in mammalian teeth (Naleway *et al*., 2015), the tendon-ligament (Lu & Thomopoulos, 2013), collagen/elastin of skin (Labroo *et al*., 2021) and chitin/protein in squid beaks (Miserez *et al*., 2008).

Such mechanical gradients are mainly driven by changes in local chemical composition or arrangements of the building blocks (Li *et al*., 2021). In this regard, the stiffening of the CPM could be related to the higher cross-linking observed in mature tomato cutin polyester (Philippe *et al*., 2016; Chatterjee *et al*., 2016) including the contribution of pCA (Reynoud *et al*., 2021). In addition, pCA might participate in the stiffening of cuticle through cross-linking by peroxidase action. Indeed, the application of exogenous peroxidase on isolated cuticle results in mechanical stiffening regardless of the developmental stages (Andrews *et al*., 2002). In cuticle, phenolic acids are present from early developmental stages (González Moreno *et al*., 2022; Reynoud *et al*., 2022) and are known targets of peroxidase. The peroxidase driven stiffening of the CPM is therefore possible, as peroxidase activity considerably increases in the epidermis during fruit ripening along with the phenolic burst (Thompson *et al*., 1998). From the isolated cuticle tensile tests, flavonoids accumulation during the fruit maturation have been associated to the increase of elastic modulus (Domínguez *et al*., 2009; Benítez *et al*., 2021). However, in-depth imaging of the CPM during tomato fruit development showed that the spatial accumulation of flavonoids at the maturation stage did not correlate with a high elastic modulus. In cuticle, a fraction of the flavonoids is readily extracted with waxes while another one is tightly embedded in the CPM (Hunt & Baker, 1980; Luque *et al*., 1995). Accordingly, our results suggest that stiffening impacts of flavonoids should rather be attributed to the solvent soluble fraction than to the CPM-associated fraction.

Besides, our recent data showed that polysaccharides embedded in the cutin matrix are subjected to macromolecular modifications during tomato fruit development (Reynoud *et al*., 2022). Indeed, the pectin to cellulose ratio within the tomato cuticles varies spatially and temporally, while the cellulose crystallinity and hemicellulose remodeling increase during fruit development. The contribution of the embedded highly esterified pectins to the mechanical properties is still unclear (Bidhendi & Geitmann, 2016) while the modification of the crystalline cellulose distribution will likely affect the in-depth mechanical properties. Likewise, pectin deposition appears essential for the assembly of cellulose microfibrils as abnormal cellulose organization was observed for an *Arabidopsis thaliana* mutant impaired in pectin biosynthesis (Du *et al*., 2020). Thus, the tight pectin-cellulose interactions observed in the CPM (Reynoud *et al*., 2022) and in the cell wall (Wang *et al*., 2015) should significantly impact the CPM mechanical properties. Accordingly, the higher structural order of both pectin and cellulose during fruit maturation might contribute to the progressive stiffening of the CPM. In the same extent, hemicelluloses were largely remodeled in the CPM during development (Reynoud *et al*., 2022). This should modify their binding to cellulose (Grantham *et al*., 2017; Jaafar *et al*., 2019) which affects microfibrils organization (Cosgrove, 2022), hence supporting a modification of the mechanical properties of CPM.

Finally, specific molecular orientations were observed in the CPM area with distinct mechanical properties (Fig. **5a**). Such correlation could be inferred by analogy to the contribution of microfibrils orientations of cellulose in the load-bearing capacity of the cell wall (Cosgrove, 2022). It should be noted that in the CPM, specific orientations of cellulose microfibrils were observed that were modified during tomato fruit development (Reynoud *et al*., 2022). Due to periclinal expansion, progressive microfibrils reorientation and straightening (Renaudin *et al*., 2017; Zhang *et al*., 2021) would results in higher elastic modulus. Indeed, macromolecular straightening is monitored through the persistence length, i.e., the distance over which molecular chain is aligned to main tangential axis (Flory & Volkenstein, 1969). An increase in the persistence length results in a more stiff rod (with a higher elastic modulus), than a more coiled or bend structure (Usov *et al*., 2015; Zdunek *et al*., 2021). The same conclusion could be drawn for pectins which harbored an ordered structures and specific orientation in the CPM during fruit maturation (Reynoud *et al*., 2022). These macromolecular modifications could account for the increase in elastic modulus over fruit development.

Altogether, the locally heterogeneous mechanical properties of CPM are finely tuned during the development of tomato fruit and are related to different local variations of chemical compositions, macromolecular arrangements and distribution. Such multiplicity of gradients will likely provide an architectural basis to fit the CPM mechanical properties with the mechanical stress imposed during the different phases of fruit expansion and ripening. This fine tuning of cuticle architecture and its mechanical properties provide new insight for plant breeding as well as the design of bioinspired functional material.

## Acknowledgements

Raman and AFM were performed at the PROBE research infrastructure, Biopolymers Interactions, Structural Biology (BIBS) facility, Nantes. OPTIR was performed at the Institut de Chimie Physique. NR was supported by Ph.D. fellowship (SeaSCAPE) granted by INRAE and the Region Pays de la Loire. This work was also supported by the ANR (Agence National de la Recherche) grant COPLAnAR (ANR-21-CE11-0035).

## Author contributions

CR, ML, DM and BB designed the research. NR, NG, JP, JM and ADO performed experiments. NR, NG, ADO, JM, ADB, ML, DM, CR and BB analyzed the data. NR, ML, DM, CR and BB wrote the paper.

## Supporting Information

**Fig. S1** In-depth mechanical properties of the cutin polymer matrix over tomato fruit development.

**Fig. S2** Mechanical properties of the central furrow of the cutin polymer matrix over tomato fruit development.

**Fig. S3** Cluster analysis of hyperspectral maps of the tomato cutin polymer matrix at 25 DPA stage.

**Fig. S4** In-depth Raman profile of the cutin polymer matrix over tomato fruit development.

**Fig. S5** Hyperspectral profiling of the central furrow and the furrow sides of the cutin polymer matrix at 20, 30 and 40 days post anthesis (DPA).

**Fig. S6** Loadings of principal component analysis of OPTIR and Raman datasets at 40 days post anthesis (DPA).

**Fig. S7** Macromolecular orientation of lipids within the central furrow compared to the furrow sides at 40 DPA.

