## Supplementary material for "Cuticle architecture and mechanical properties: a functional relationship delineated through correlated multimodal imaging": Sup information

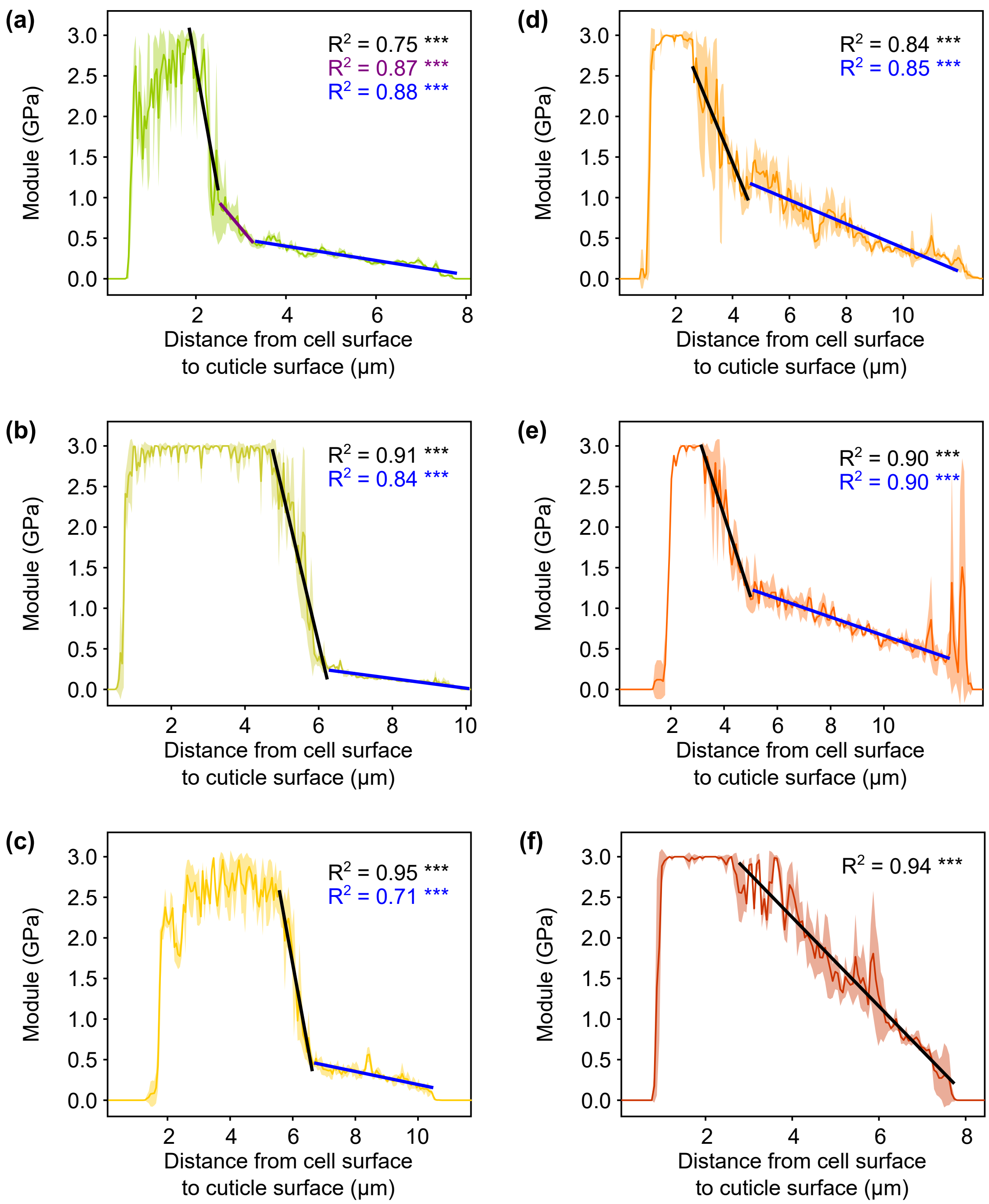


**Fig. S1** In-depth mechanical properties of the cutin polymer matrix over tomato fruit development. Radial profile from the epidermal cell surface to the cuticle surface for (a) 15, (b) 20, (c) 25, (d) 30, (e) 35 and (f) 40 days post anthesis (DPA). Each profile consists of 160 to 235 equally interspersed points, according to the in-depth cuticle thickness, with each point being a mean of 8 measurements. Two maps per developmental stage were acquired with 7 sampling lines per stage. Results for one representative map per stage are displayed. Data are expressed as mean (solid lines) ± SD (shadowed area). Linear regressions were performed on the decreasing slope of the elastic modulus with three regression for 15 DPA, two regressions for 20 to 35 DPA and one regression for 40 DPA. The quality of the fit are represented by the squared-R values.

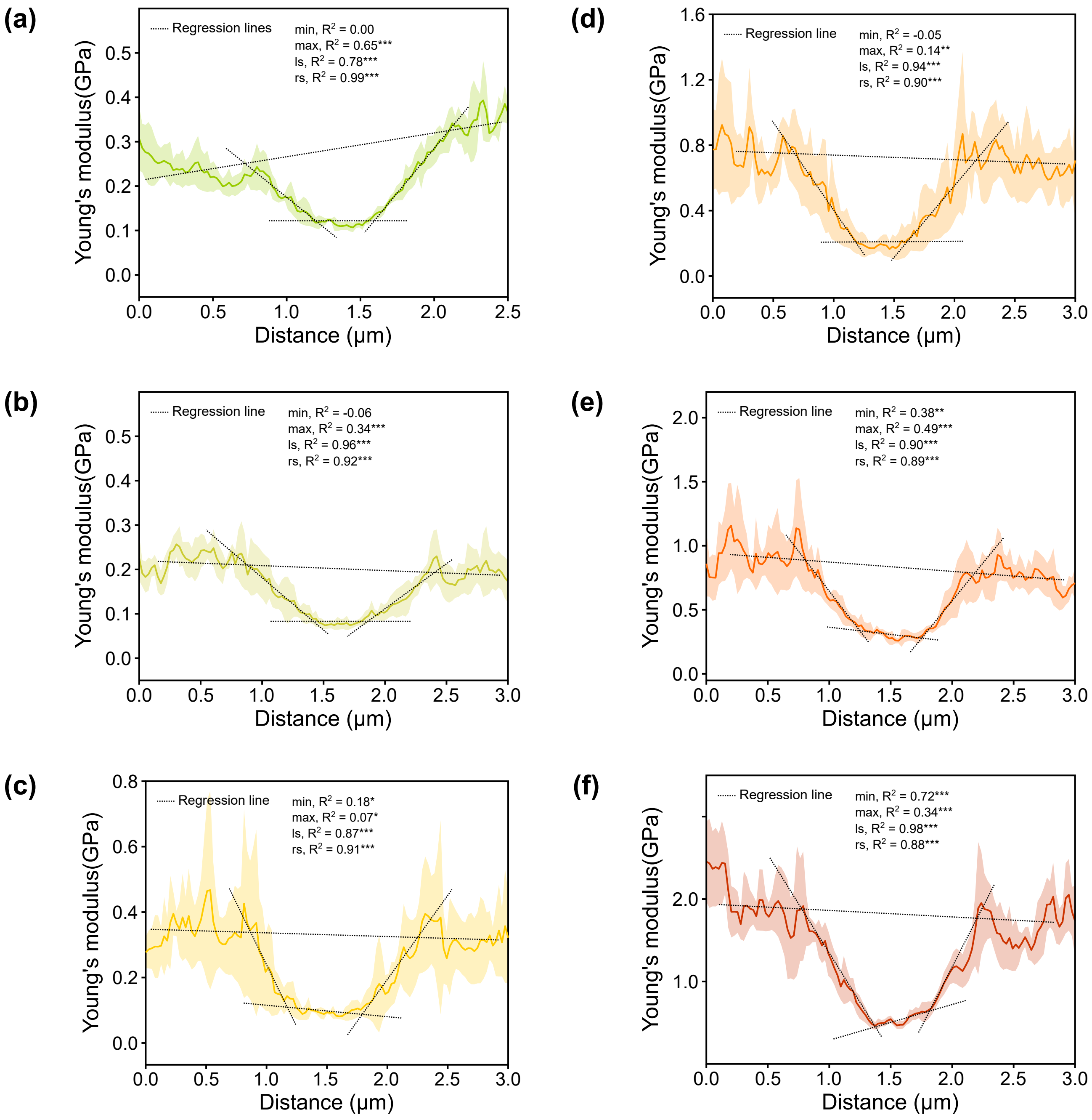


**Fig. S2** Mechanical properties of the central furrow of the CPM over tomato fruit development. Transverse profile of the central furrow for (a) 15, (b) 20, (c) 25, (d) 30, (e) 35 and (f) 40 days post anthesis (DPA). Ten profiles were sampled per developmental stage with each profile consisting of 120 equally interspersed points over 3µm; each point being a mean of 8 measurements. Two maps per developmental stage were acquired, results for one representative map per stage are displayed. Data are expressed as mean (solid lines) ± SD (shadowed area). Linear regressions were performed on the minimum (min) and maximum (max) and both sides of the slope (left side, ls and right side, rs) of the elastic modulus. The quality of the fit are represented by the squared-R values.

(a)

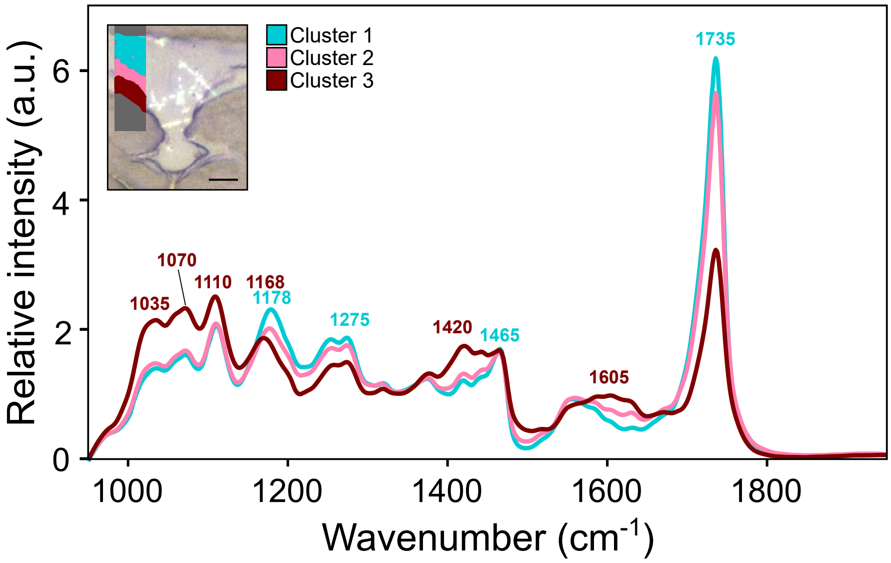


(b)

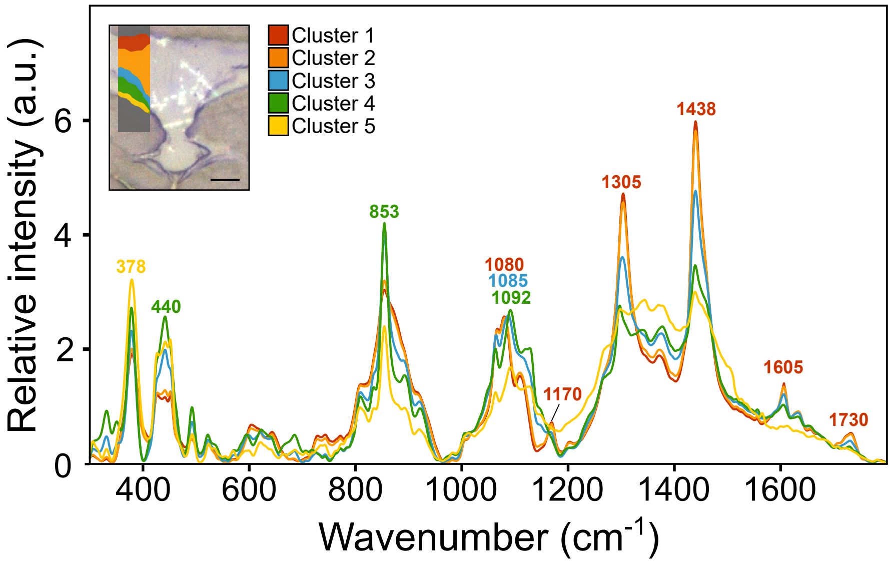


**Fig. S3** Cluster analysis of hyperspectral maps of the tomato CPM at 25 DPA stage. K-means. Insets represents K-means clustering, performed according to Reynoud et al., (2022), with charts displaying mean spectrum per cluster. The number of clusters for each dataset was tested from 2 to 10 clusters. (a) K-means clustering of OPTIR maps with three clusters appeared more appropriate. Mean spectra for cluster 1 and 2 are similar and display bands typical of lipids, i.e., 1178 $cm^{-1}$ assigned to ν as (C-O-C), 1465 $cm^{-1}$ assigned to $\delta(CH_2)$ and 1735 $cm^{-1}$ assigned to the ν (C=O) (Skolik et al., 2019), hence highlights cutinized area. Cluster 3 mean spectrum displays mainly polysaccharides fingerprints, e.g., 1035, 1070 and 1110 attributed to ν(CO, CC, CCO) (Chylińska et al., 2016), and account for cell-wall areas. (b) K-means clustering of Raman maps with 5 clusters appeared more appropriate with each cluster being distinguished in pectin (855 $cm^{-1}$), crystalline cellulose (379 $cm^{-1}$), cutin (1305 and 1440 $cm^{-1}$) and phenolics (1170 and1605 $cm^{-1}$) content. Band assignment has been done according to Reynoud et al., (2022). Scale bar = 5µm.

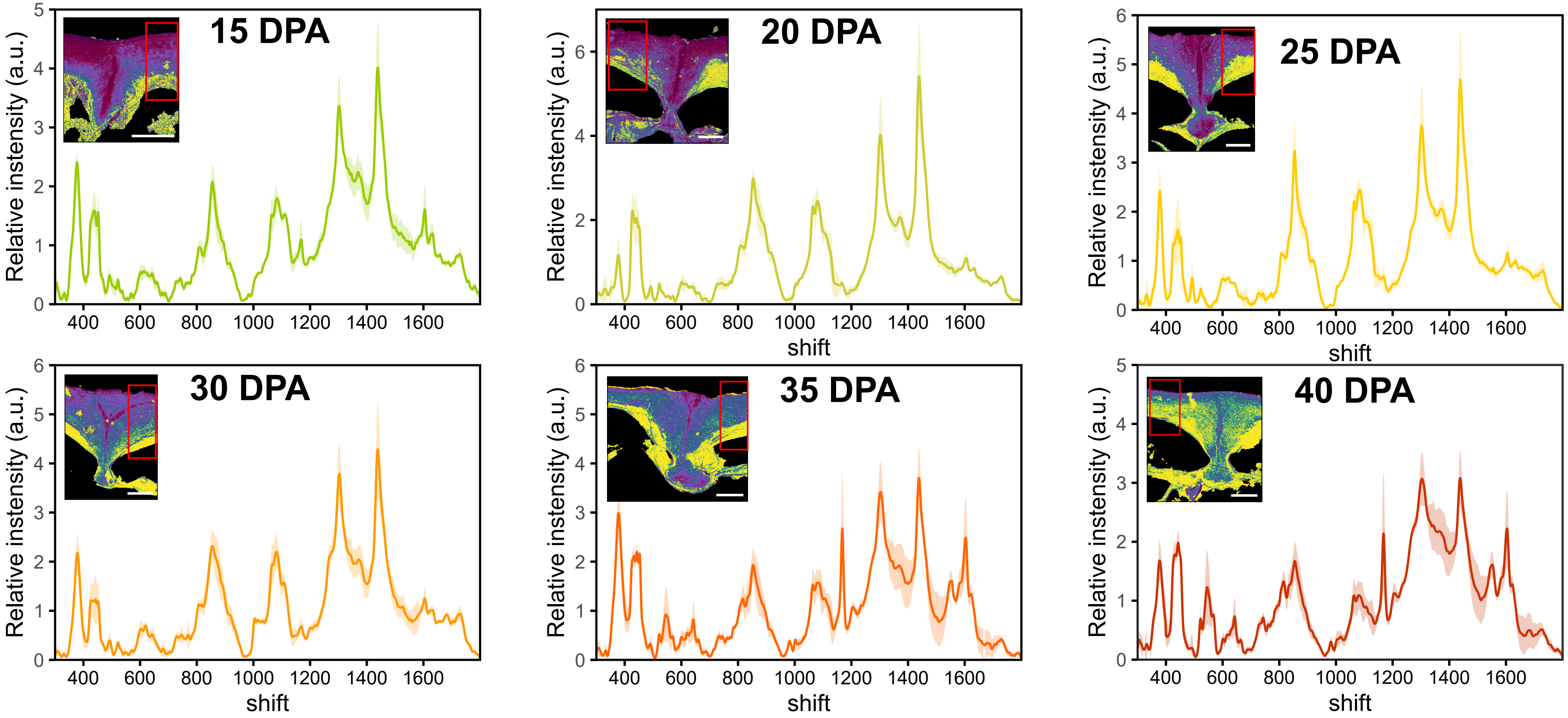


**Fig. S4** In-depth Raman profile of the CPM over tomato fruit development. Inset represent the elastic modulus map (Fig. 2) with the red rectangle indicating the area of the CPM sampled for Raman profiling. Scale bar = 5μm. Data are expressed as mean (solid lines) ± SD (shadowed area) spectra of the area sampled. Area-normalization was performed before calculating the mean spectra.

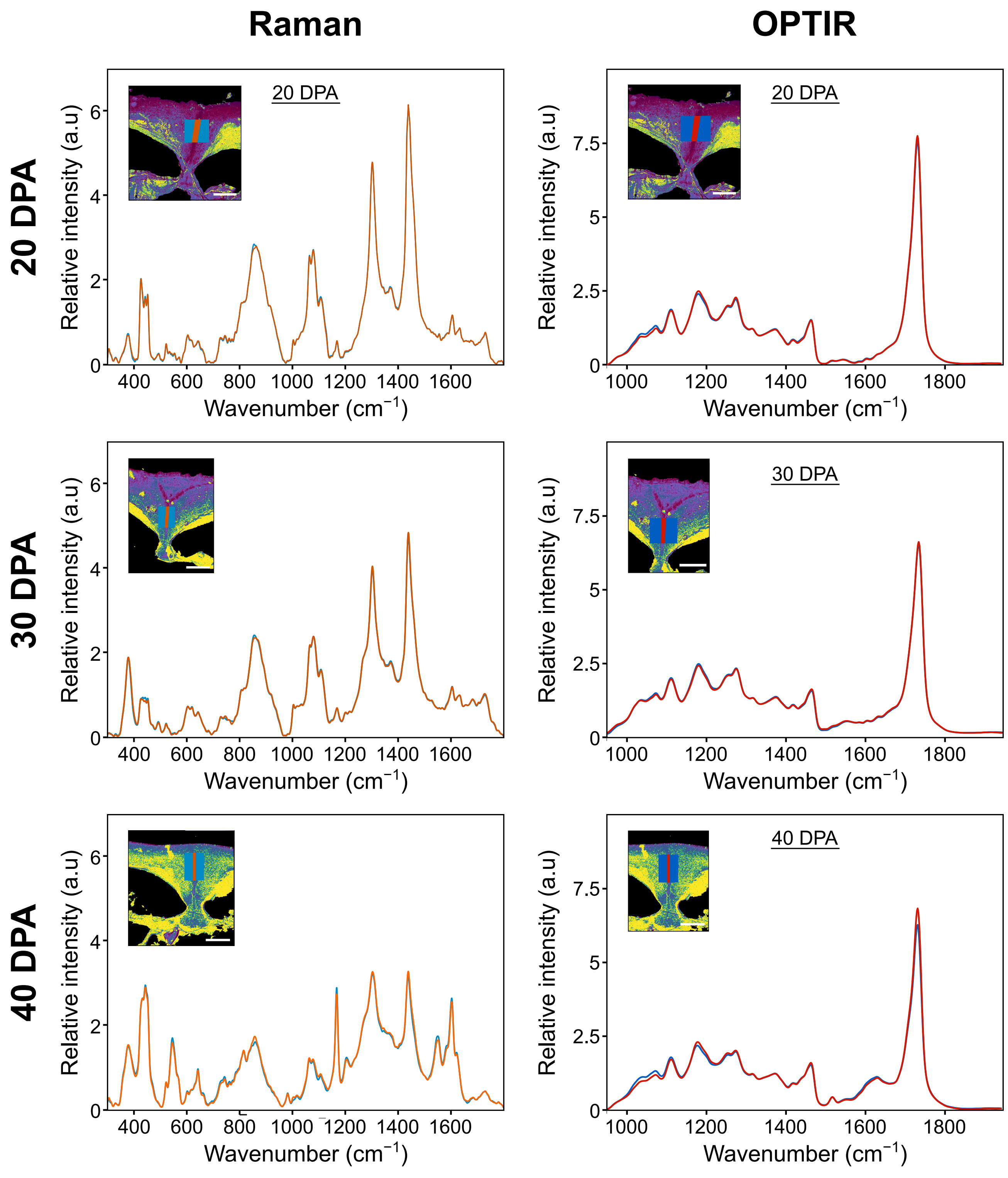


**Fig. S5** Hyperspectral profiling of the central furrow and the furrow sides of the CPM at 20, 30 and 40 days post anthesis (DPA). Insets represent the elastic modulus map (Fig. 2) with the solid rectangle indicating the area of the CPM sampled and correspond to the mean Raman spectra displayed. Area-normalization was performed before calculating the mean spectra. Scale bar = 5μm.

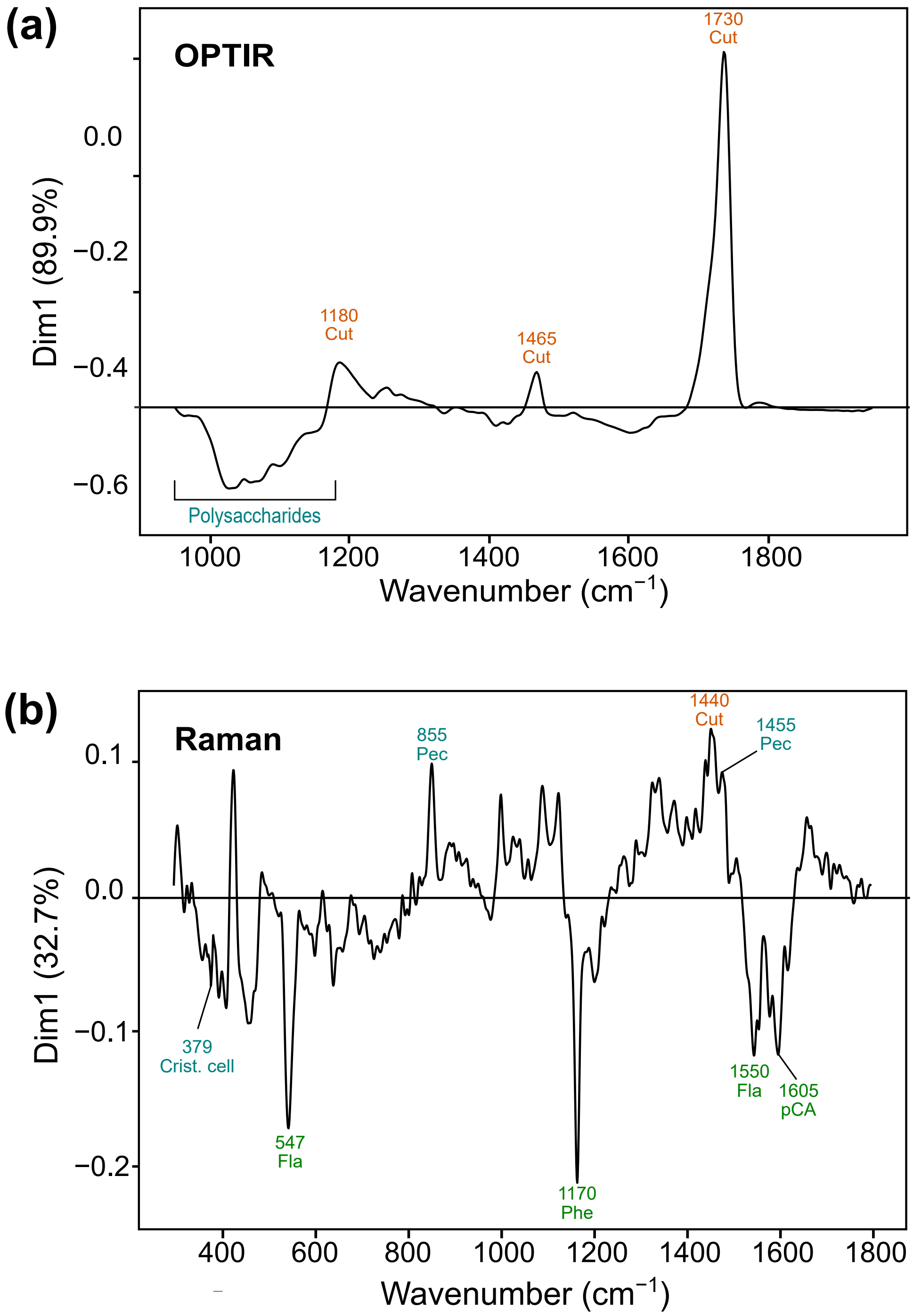


**Fig. S6** Loadings of principal component analysis (PCA) of OPTIR (up) and Raman (down) dataset at 40 DPA. The loadings refer to the Fig. 5a. Cut, cutin; Crist. cell, crystalline cellulose; Fla, flavonoids, Pec, pectin; Phe, phenolics; pCA, p-coumaric acid.

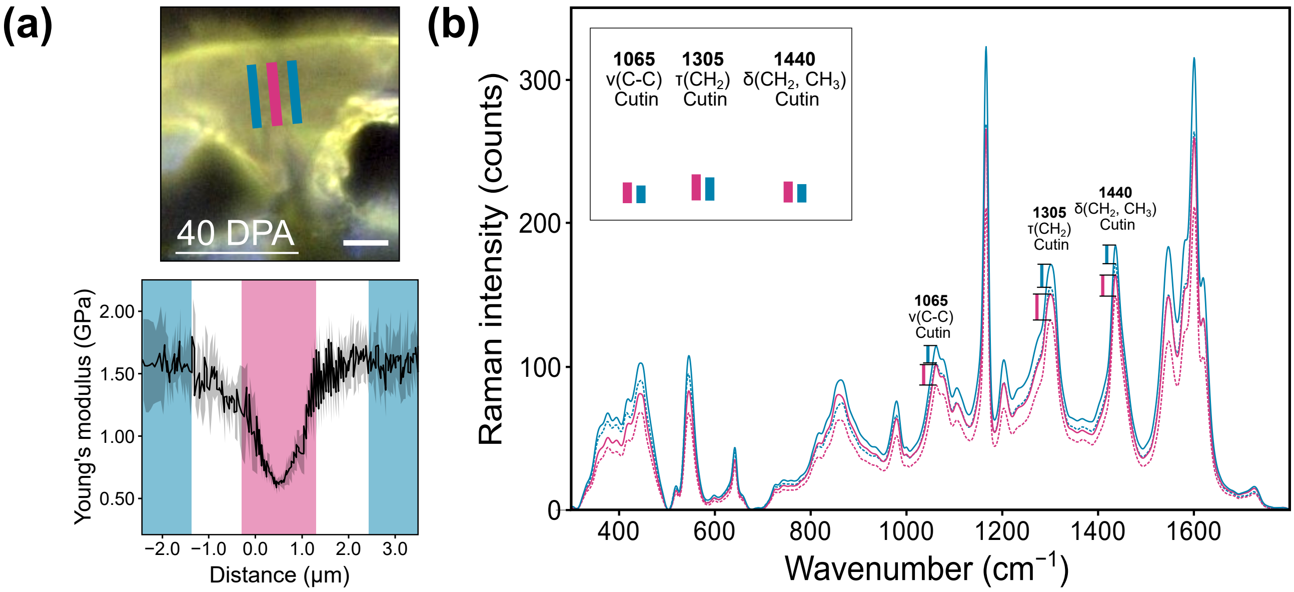


**Fig. S7** Macromolecular orientation of lipids within the central furrow compared to the furrow sides at 40 DPA. The laser polarization direction was modified from parallel (0°) to orthogonal (90°) to the cuticle surface. For each polarization, two maps were acquired. The light image (upper left) represents the area of the cuticle that were sampled to assess the molecular orientations within the central furrow (pink) and the furrow sides (blue) with the corresponding elastic modulus (lower left). Scale bar = 5µm. Data are expressed as mean (solid lines) ± SD (shadowed area). For both central furrow and furrow sides, mean Raman spectra for each polarization were calculated (right). The parallel (solid lines) to orthogonal (dashed lines) ratio of intensity was illustrated by vertical bars for three bands related to lipids: two bands attributed to $CH_2$ deformations, i.e., τ($CH_2$) at 1305 cm$^{-1}$ and δ($CH_2$,$CH_3$) at 1440 cm$^{-1}$ and one band assigned to ν(C-C) vibration at 1065 cm$^{-1}$. Insets represents the different intensity ratio lined up (bars) for proper comparison between cuticle areas.
